# Rabphilin 3A binds the N-peptide of SNAP-25 to promote SNARE complex assembly in exocytosis

**DOI:** 10.1101/2022.05.17.492271

**Authors:** Tianzhi Li, Qiqi Cheng, Shen Wang, Cong Ma

## Abstract

Exocytosis of secretory vesicles requires the soluble N-ethylmaleimide-sensitive factor attachment protein receptor (SNARE) proteins and small GTPase Rabs. As a Rab3/Rab27 effector protein on secretory vesicles, Rabphilin 3A was implicated to interact with SNAP-25 to regulate vesicle exocytosis in neurons and neuroendocrine cells, yet the underlying mechanism remains unclear. In this study, we have characterized the physiologically relevant binding sites between Rabphilin 3A and SNAP-25. We found that an intramolecular interplay between the N-terminal Rab-binding domain and C-terminal C_2_AB domain enables Rabphilin 3A to strongly bind the SNAP-25 N-peptide region via its C_2_B bottom α-helix. Disruption of this interaction significantly impaired docking and fusion of vesicles with the plasma membrane in PC12 cells. In addition, we found that this interaction allows Rabphilin 3A to accelerate SNARE complex assembly. Furthermore, we revealed that this interaction accelerates SNARE complex assembly via inducing a conformational switch from random coils to α-helical structure in the SNAP-25 SNARE motif. Altogether, our data suggest that promotion of SNARE complex assembly by binding of the C_2_B bottom α-helix of Rabphilin 3A to the N-peptide of SNAP-25 underlies a pre-fusion function of Rabphilin 3A in vesicle exocytosis.

## Introduction

Secretion of neurotransmitters and hormones mediated by Ca^2+^-regulated exocytosis in neurons and neuroendocrine cells is achieved by a series of intracellular membrane trafficking steps, including recruitment, docking, priming and fusion of secretory vesicles with the plasma membrane (Rettig and Neher, 2002; Südhof and Rizo, 2011; Jahn and Fasshauer, 2012). The core machinery governing the process involves the SNARE proteins synaptobrevin-2 (Syb2) on secretory vesicles, and syntaxin-1 (Syx1) and SNAP-25 (SN25) on the plasma membrane, which form the four helical bundle structure called the SNARE complex to bring the two membranes into close proximity and eventually drive membrane fusion (Jahn and Scheller, 2006; Sutton et al., 1998; Weber et al., 1998). Small GTPase Rabs are essential regulators in the secretory pathway, which have specific role in defining the identity of subcellular membranes (Stenmark, 2009). Rab3A and Rab27A are abundantly expressed in neurons and neuroendocrine cells and associate with the membrane of secretory vesicles in a GTP-bound form to regulate vesicle exocytosis (Takai et al., 1996; Geppert and Südhof, 1998; Fukuda, 2005). Rabphilin 3A (Rph3A) was originally identified as a GTP-Rab3A-binding protein and found to regulate SNARE-dependent exocytosis (Shirataki et al., 1992; Tsuboi and Fukuda, 2005; Deak et al., 2006). Recent studies have found that loss of Rph3A is associated with Alzheimer’s and Huntington’s disease, suggesting a crucial role of Rph3A in synaptic function (Smith et al., 2005; Smith et al., 2007; Tan et al., 2014).

Rph3A contains an N-terminal Rab-binding domain (RBD) that associates with secretory vesicles via binding to GTP-bound Rab3A or Rab27A (Li et al., 1994; Mizoguchi et al., 1994; Fukuda et al., 2004), a proline-rich linker region (PRL) bearing multiple phosphorylation sites (Foletti et al., 2001), and two C-terminal C_2_ domains (termed C_2_A and C_2_B, respectively) that interact with phospholipids and the SNAREs (Yamaguchi et al., 1993; Chung et al., 1998; Tsuboi and Fukuda, 2005) (***Figure 1A***). In particular, increasing evidence showed that the C_2_B domain of Rph3A binds SN25 to regulate exocytosis of dense-core vesicles (DCVs) in PC12 cells and fine-tune re-priming of synaptic vesicles in hippocampal neurons (Tsuboi and Fukuda, 2005; Deak et al., 2006), but the underlying mechanism was unclear. It was previously reported that Rph3A binds SN25 via the β3-β4 polybasic region of the C_2_B domain (Tsuboi et al., 2007), while later structural results identified that Rph3A interacts with SN25 via the α-helix at the bottom face of the C_2_B domain (Deak et al., 2006; Ferrer-Orta et al., 2017). In addition, according to the crystal structure of the Rph3A-C_2_B–SN25 complex, three distinct sites in SN25 were found to participate in binding to Rph3A-C_2_B, two of which located in the middle region of the SNARE motif of SN25, and one resided at the N-terminus of SN25 (Ferrer-Orta et al., 2017). However, it remains elusive which binding mode of the Rph3A-C_2_B–SN25 interaction is physiologically relevant. Importantly, little is known how this interaction regulates vesicle exocytosis.

**Figure 1.**
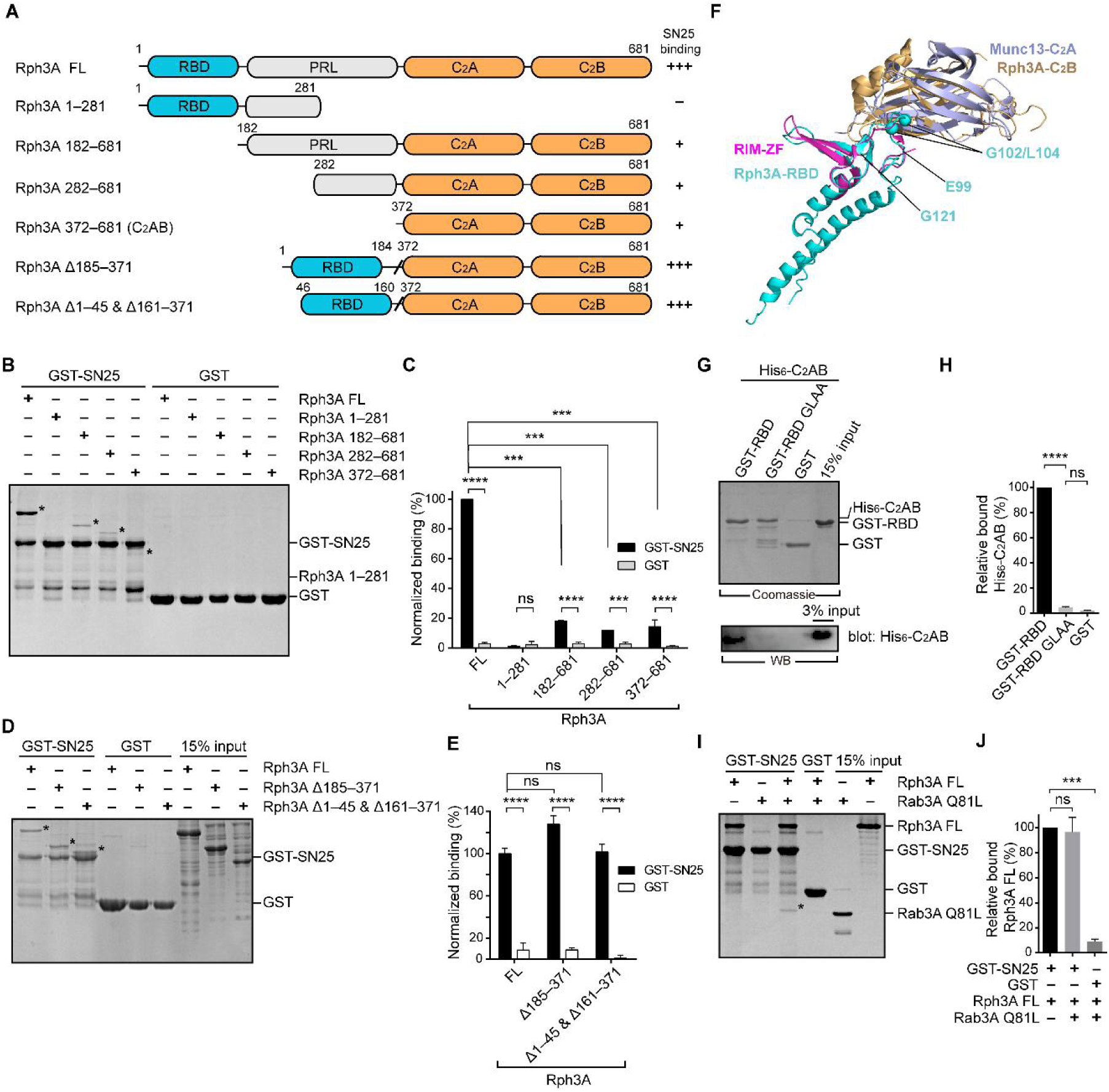
The intramolecular interplay of Rph3A enables strong binding to SN25. (A) Schematic diagram showing domain organization and variant fragments of Rph3A. The relative SN25 binding affinity (–, +, +++) derived from binding experiments is indicated on the right. (B and C) Binding of Rph3A FL or the fragments to GST-SN25 measured by GST pull-down assay (B) and quantification of the Rph3A binding (C). Asterisks in B show bands of bound Rph3A FL or fragment proteins. (D and E) Binding of Rph3A FL or deletion mutations (Δ185–371, Δ1–45 &Δ161–371) to GST-SN25 measured by GST pull-down assay (D) and quantification of the Rph3A binding (E). Asterisks in D show bands of bound Rph3A FL or fragment proteins. (F) Structure alignment of Rph3A with RIM-ZF–Munc13-C2A complex (PDB entry: 2CJS). The Rph3A RBD (PDB entry: 1ZBD) and C2B (PDB entry: 3RPB) domain were aligned and superposed with RIM-ZF, Munc13-C2A, respectively. The G102/L104 site was mapped to Rph3A-RBD structure. (G and H) Binding of His6-tagged Rph3A-C2AB to GST-Rph3A-RBD (40–170) or its G102A/L104A mutant (GLAA) measured by GST pull-down assay (G) and quantification of the bound His6-C2AB (H). The top panel shows a Coomassie blue stained gel of the GST-proteins to illustrate that similar amounts of protein were employed. Bound His6-C2AB proteins were analyzed by immunoblotting with anti-His6 antibody (bottom). (I and J) Binding of Rph3A FL to the GST-SN25 in the presence of Rab3A Q81L measured by GST pull-down assay (I) and quantification of the bound Rph3A (J). Asterisk in I shows the band of bound Rab3A Q81L. Data are processed by Image J (NIH) and presented as the mean ± SEM (n = 3), technical replicates. Statistical significance and P values were determined by one-way analysis of variance (ANOVA). ***P < 0.001; ****P < 0.0001; ns: not significant.

In this study, we have systematically characterized the interaction between Rph3A and SN25 using full-length form of Rph3A. We found that the N-peptide of SN25 (residues 1–10) mediates the Rph3A–SN25 interaction in a manner that requires an intramolecular interplay between the N-terminal RBD domain and C-terminal C_2_AB domain of Rph3A. In addition, we identified that the bottom α-helix rather than the β3-β4 polybasic region in the C_2_B domain of Rph3A mediates SN25 interaction. Deletion of the N-peptide of SN25 or disruption of the C_2_B bottom α-helix of Rph3A significantly reduced the Rph3A–SN25 interaction and impaired docking and fusion of DCVs with the plasma membrane in PC12 cells, revealing the physiological relevance of such Rph3A–SN25 interaction. Furthermore, we found a stimulatory role of Rph3A in SNARE complex assembly dependent on the Rph3A–SN25 interaction. This stimulatory effect of Rph3A arises because Rph3A induces a conformational change of the SN25 SNARE motif from random coils to α-helical structure.

## Results

### An intramolecular interplay of Rph3A FL contributes to SN25 interaction

Given the difficulties in obtaining full-length Rph3A (Rph3A FL, residues 1–681) proteins *in vitro*, most previous studies alternatively used various Rph3A fragments to map the potential binding sites between Rph3A and SN25 (Deak et al., 2006; Ferrer-Orta et al., 2017). Here, we have successfully obtained Rph3A FL with desirable yield in *E. coli* by using a pEXP5-NT/TOPO expression system. Rph3A FL displayed a strong tendency to form monomer in solution, as detected by size-exclusion chromatograph and analytical ultracentrifugation (***Figure 1-figure supplement 1***).

We first examined the binding between Rph3A and SN25 (full-length, 1–206) by using GST pull-down experiments. We have designed and obtained a series of truncations that selectively comprise one or several domains of Rph3A, i.e., Rph3A (1–281), Rph3A (182–681), Rph3A (282–681) and Rph3A (372–681) (***Figure 1A***). As controls, GST alone displayed no binding to Rph3A FL and the truncations. In contrast, GST-SN25 showed significant interaction with Rph3A (182–681), Rph3A (282–681) and Rph3A (372–681), but failed to bind Rph3A (1–281) (***Figure 1B, C***). Consistent with previous observations, these data indicate that the C-terminal C_2_AB of Rph3A contributes predominately to SN25 interaction. Intriguingly, GST-SN25 showed much more stronger binding capacity to Rph3A FL than to the above truncations (***Figure 1B, C***). As Rph3A (1–281) did not bind SN25, the only explanation is that the N-terminal region of Rph3A cooperates with the C-terminal C_2_AB to enhance SN25 binding.

To determine which part in the N-terminal region of Rph3A is required for the enhanced SN25 binding, we generated two Rph3A variants, one selectively lacked the middle PRL, i.e., Rph3A (Δ185–371), the other additionally removed the N- and C-terminal residues upstream and downstream of the RBD, i.e., Rph3A (Δ1–45 & Δ161–371) (***Figure 1A***). Both variants bound to GST-SN25 as effectively as Rph3A FL (***Figure 1D, E***), revealing that the RBD is important for the enhanced SN25 binding. As Rph3A FL prefers a monomeric state in solution and that the RBD alone displayed no detectable binding to GST-SN25 as shown in ***Figure 1B, C***, we suspect that an intramolecular interplay between the RBD and the C_2_AB may enhance Rph3A–SN25 binding. Indeed, we observed that GST-tagged Rph3A-RBD (40–170) but not GST alone bound to His_6_-tagged Rph3A-C_2_AB (372–681), as detected by pull-down assay combined with immunoblotting (***Figure 1G, H***). Next, we sought to explore the binding sites in the RBD that mediate binding to the C_2_AB. We used the crystal structure of the zinc-finger (ZF) of RIM-1 bound to the C_2_A domain of Munc13-1 as a guide (Lu et al., 2006), as Rph3A-RBD and Rph3A-C_2_AB similarly contain zinc-finger (ZF) sequence and C_2_-domain sequence, respectively. Superimposition of the structures of Rph3A-RBD and Rph3A-C_2_B with the structure of the RIM-ZF–Munc13-C_2_A complex points to a region (residues 99–121) in the RBD (***Figure 1F***). Upon screening, we found that mutation of G102/L104 (G102A/L104A, referred to as GLAA) abolished binding of Rph3A-RBD to Rph3A-C_2_AB, as detected by immunoblotting (***Figure 1G, H***). To confirm this, we also introduced the GLAA mutation to Rph3A (Δ185–371) and found that it severely impaired GST-SN25 binding (***Figure 1-figure supplement 2A, B***). Altogether, these data suggest a new binding mode between Rph3A and SN25 which relies on an intramolecular interplay of the RBD with C_2_AB in Rph3A.

The RBD of Rph3A can bind Rab3A (Q81L, GTP-bound form) as well (Stahl et al., 1996; Ostermeier and Brunger, 1999). Based on the crystal structure of the Rph3A-RBD–Rab3A complex (Ostermeier and Brunger, 1999), the binding sites for Rab3A position away from residues G102/L104 in the RBD (***Figure 1-figure supplement 2C***), suggesting that the intramolecular interplay of the RBD with C_2_AB might be compatible with Rab3A interaction. Indeed, pull-down experiments showed that the involvement of Rab3A didn’t influence the strong interaction between GST-SN25 and Rph3A FL mediated by the intramolecular interplay of the RBD with C_2_AB (***Figure 1I, J***). Instead, Rab3A showed significant binding to the Rph3A FL–SN25 complex (***Figure 1I, J***), implying that Rph3A is able to serve as a connector to bridge vesicle-bound Rab3A and plasma membrane-bound SN25.

### The N-peptide of SN25 mediates Rph3A FL interaction

SN25 contains two SNARE motifs, SNARE motif 1 and 2, connected by a linker region, namely SN1, SN2, and LR, respectively. SN25 also comprises an N-terminal peptide preceding SN1 (***Figure 2A***). Previous structural analysis using the C_2_B fragment of Rph3A showed that both the middle part of SN1 and the N-peptide of SN25 are involved in Rph3A-C_2_B interaction (Ferrer-Orta et al., 2017). According to the binding mode of Rph3A with SN25 as described above (***Figure 1***), it is necessary to pinpoint the binding sites on SN25 by using Rph3A FL rather than the C_2_B fragment. To this aim, we designed and obtained a series of SN25 fragments that selectively contains multiple domains, e.g., SN25 (1–82), SN25 (1–140), SN25 (141–206), SN25 (83–206), and SN25 (83–140), and compared their binding abilities with Rph3A FL (***Figure 2A***). Among these fragments, SN25 (1–82) and SN25 (1–140) significantly bound to Rph3A FL, as effectively as SN25 FL (***Figure 2B, C***). In contrast, GST alone and the other SN25 fragments failed to bind Rph3A FL (***Figure 2B, C***). Hence, these data suggest that Rph3A FL binds to the N-terminal region of SN25 comprising the SN1 and the N-peptide.

**Figure 2.**
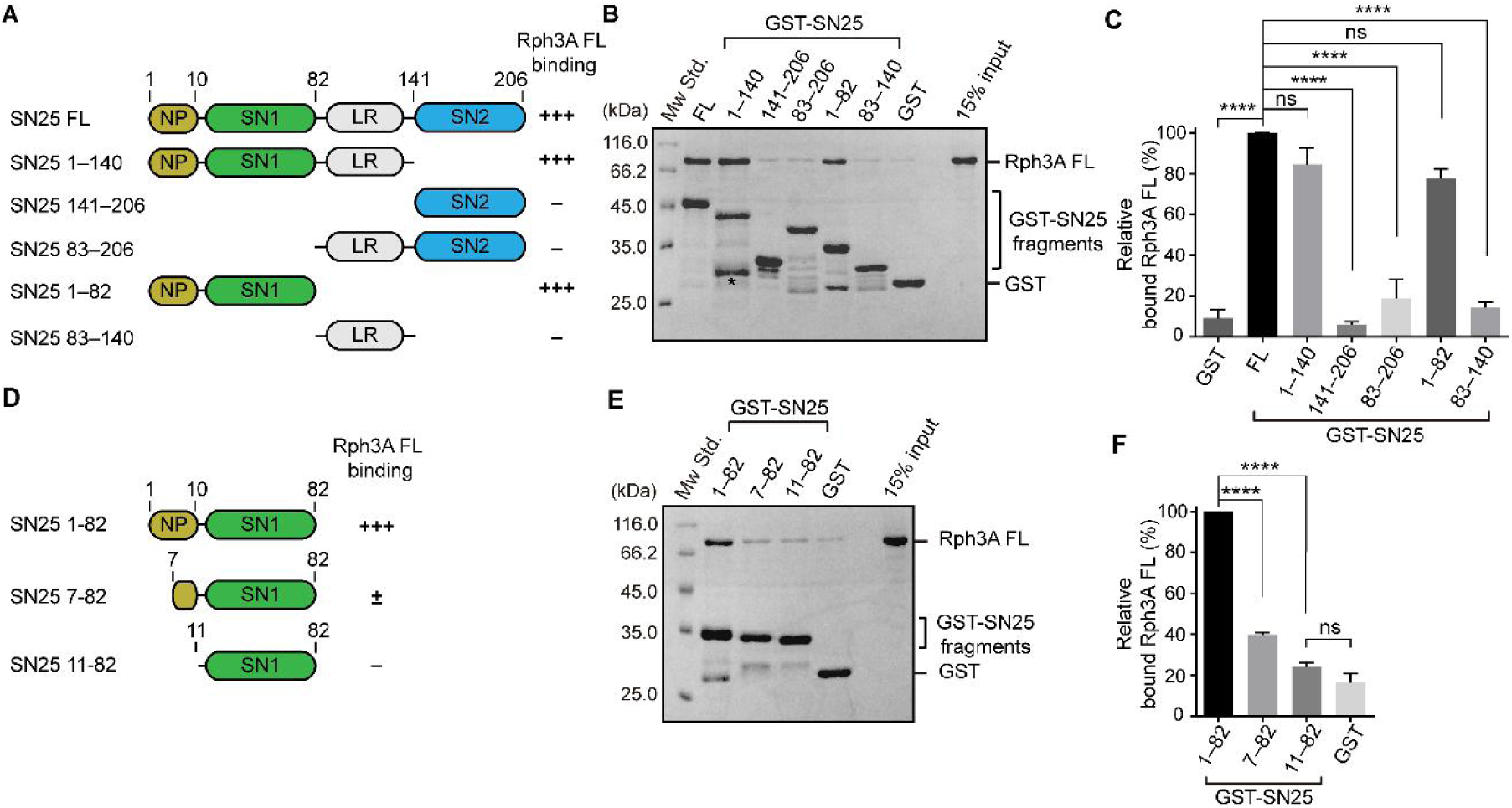
Characterization of the Rph3A-binding sites in SN25. (A) Schematic diagram showing domain organization and variant fragments of SN25. NP, N-peptide of SN25 (residues, 1–10); SN1, SNARE motif 1; LR, linker region; SN2, SNARE motif 2. The relative Rph3A FL binding affinity (–, +++) derived from binding experiments is indicated on the right. (B and C) Binding of Rph3A FL to GST-SN25 FL or fragments measured by GST pull-down assay (B) and quantification of the bound Rph3A FL (C). Asterisk in B shows the band of degraded GST-SN25 (1–140) proteins. (D) Schematic diagram showing N-peptide deletion mutants of SN25 (1–82). The relative Rph3A binding affinity (–, ±, +++) derived from binding experiments is indicated on the right. (E and F) Binding of Rph3A FL to GST-SN25 (1–82) or N-peptide deletions (7–82 and 11–82) measured by GST pull-down assay (E) and quantification of the bound Rph3A (F). Data are processed by Image J (NIH) and presented as the mean ± SEM (n = 3), technical replicates. Statistical significance and P values were determined by one-way analysis of variance (ANOVA). ****P < 0.0001; ns: not significant.

To further identify the sequence of SN25 responsible for Rph3A FL interaction, we generated two truncations on SN25 (1–82), i.e., SN25 (7–82) and SN25 (11–82), with the N-peptide residues differentially deleted (***Figure 2D***). To our surprise, in contrast to the strong interaction of GST-SN25 (1–82) with Rph3A FL, GST-SN25 (7–82) strongly impaired Rph3A FL binding, and GST-SN25 (11–82) failed to bind Rph3A FL (***Figure 2E, F***), showing that the N-peptide sequence (^1^MAEDADMRNE^10^) is essential for mediating binding of SN25 to Rph3A FL. However, previous structural and biochemical analysis using the C_2_B fragment of Rph3A suggested the middle portion of SN1 (E38/D41/R45) as the primary binding sites for Rph3A (Ferrer-Orta et al., 2017). To figure this out, we made a SN25 (1–82) mutant which carries E38A/D41A/R45A (EDR) and examined its interaction with Rph3A FL. Similar to SN25 (1–82), the EDR mutant retained strong binding to Rph3A FL (***Figure 2–figure supplement 1***), suggesting that the N-peptide rather than the SN1 mediates Rph3A FL interaction. Taken together, these data indicate that the binding mode between Rph3A FL and SN25 is distinct from the mode between Rph3A-C_2_B and SN25, in line with the results shown in ***Figure 1***. In addition, sequence alignment showed that the negatively charged N-peptide sequence of SN25 is conserved among different species (***Figure 2–figure supplement 2***), implying that SN25 N-peptide-mediated Rph3A FL interaction might be crucial for vesicle exocytosis.

### The N-peptide of SN25 is essential for vesicle docking and fusion in PC12 cells

It was previously reported that Rph3A regulates exocytosis of DCV in neuroendocrine cells (Tsuboi and Fukuda, 2005). We aimed to explore whether the N-peptide of SN25 is required for DCV docking and fusion by using knockdown-rescue approach in cultured PC12 cells. Endogenous SN25 expression was strongly suppressed by virally delivered shRNAs (see Materials and Methods) as previously described (Cahill et al., 2006; Zhou et al., 2019; Wang et al., 2020) (***Figure 3A***). In parallel, neuropeptide Y (NPY)-td-mOrange2 was applied to visualize DCVs. First, we monitored the dynamics of NPY-td-mOrange2-labeled DCVs near the plasma membrane by TIRF microscopy and counted the number of plasma membrane-associated DCVs in resting PC12 cells. As expected, SN25 knockdown cells showed significantly reduced number of plasma membrane-docked DCVs, in comparison with control cells (***Figure 3B, C***). Expression of SN25 (full-length, 1–206) restored the number of docked DCVs, whereas expression of the SN25 mutant (11–206) that lacks the N-peptide exhibited strongly impaired ability to restore the docked DCVs in PC12 cells (***Figure 3B, C***), revealing that the N-peptide of SN25 is required for DCV docking. Next, we analyzed the NPY-td-mOrange2 release events in PC12 cells under high KCl stimulation. The total number of release events in SN25 knockdown cells was significantly fewer than that in control cells (***Figure 3D, E***). Consistent with the docking phenotypes, expression of SN25 rescued the DCV release events, but expression of the SN25 mutant (11–206) failed to do so (***Figure 3D, E***). Together, these results indicate that SN25 N-peptide-mediated Rph3A interaction is required for DCV docking and fusion of exocytosis in PC12 cells.

**Figure 3.**
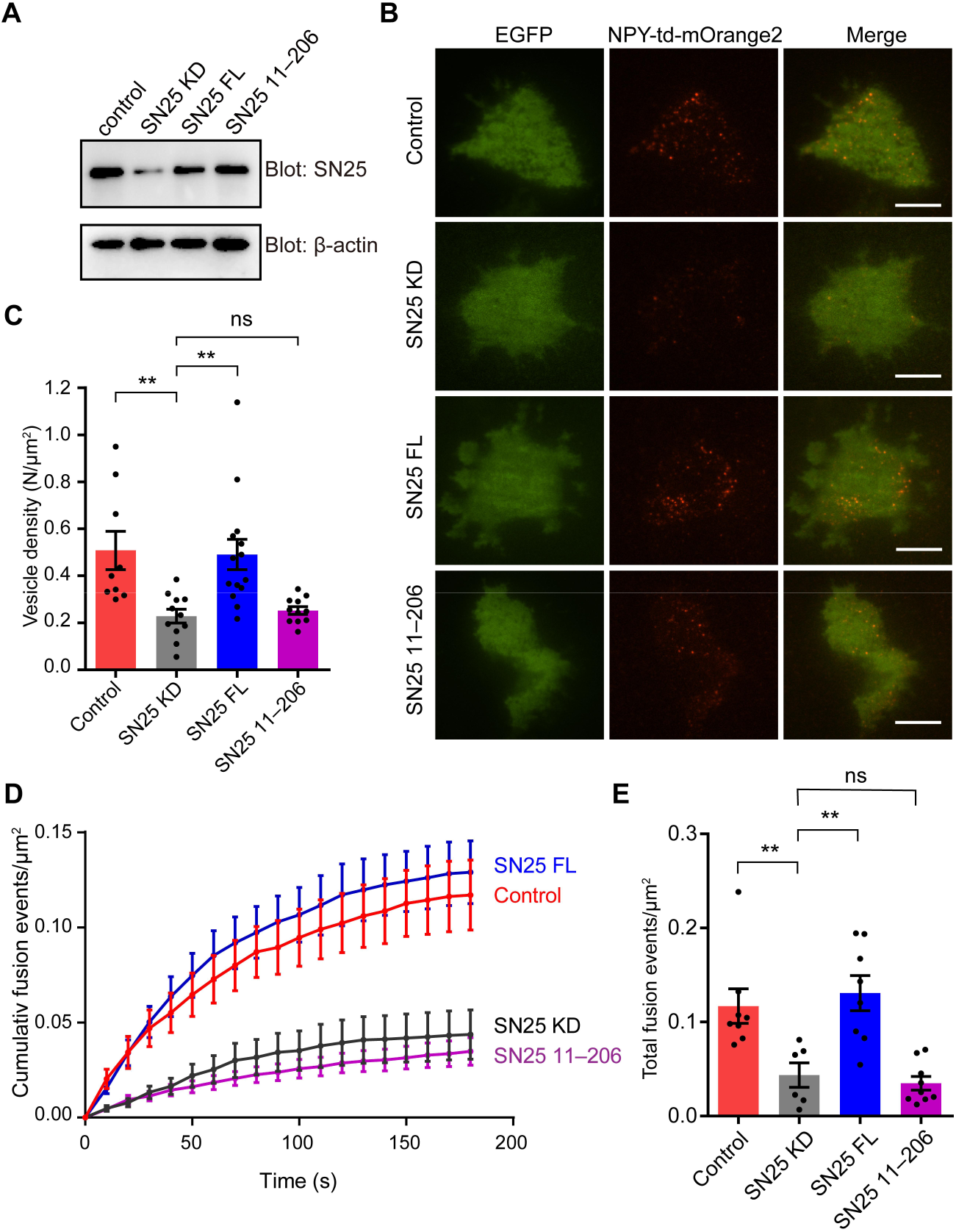
Importance of the N-peptide SN25 for DCV exocytosis in PC12 cells. (A) Immunoblotting assay to determine the SN25 expression level in control, SN25 KD, SN25 FL or SN25 (11–206) rescued PC12 cells. Expression of SN25 and β-actin in PC12 cells was analyzed by 15% SDS-PAGE followed by immunoblotting with anti-SN25 antibody, and anti β-actin antibody, respectively. (B) Representative TIRF images of control, SN25 KD, SN25 FL or SN25 (11–206) rescued PC12 cells expressing indicator EGFP, and DCV content (NPY-td-mOrange2) are shown. Scale bars, 10 μm. (C) The density of docked vesicles was determined by counting the NPY-td-mOrange2 labeled vesicles in each image (n ≥ 9 cells in each). (D) NPY-td-mOrange2 release events detected by TIRF microscopy during sustaining high K+ stimulation. The curves indicate the cumulative number of fusion events per µm2 in each cell. (E) Quantification of the results at 180 s in the experiments of (D). Data are presented as mean ± SEM; (n ≥ 6 cells in each). Statistical significance and P values were determined by one-way analysis of variance (ANOVA). **P< 0.01; ns, not significant.

### The bottom α-helix of the C_2_B domain in Rph3A FL mediates interaction with SN25

The bottom α-helix (interface I) and the β3-β4 polybasic region (interface II) in the C_2_B domain of Rph3A (***Figure 4A***), both having positively charged lysine residues, were implicated to mediate SN25 interaction (Tsuboi et al., 2007; Ferrer-Orta et al., 2017). Here, in the context of Rph3A FL, we sought to figure out which interface is physiologically relevant and represents the actual binding site for SN25. We first examined the interactions between the two interfaces and SN25 (1–82) *in vitro*. Accordingly, we designed two mutations in Rph3A FL, i.e., the K651A/K656A/K663A mutation (termed K3) in interface I and the K590Q/K591Q/K593Q/K595Q mutation (termed K4) in interface II (***Figure 4A***). Using GST pull-down assay, we found that K4 showed slightly reduced binding to SN25 (1–82) compared to Rph3A FL, whereas K3 displayed remarkably impaired interaction with SN25 (1–82) (***Figure 4B, C***). Similar results were found when the mutations were introduced to SN25 (1–140) (***Figure 4****–**figure supplement 1***). These data indicate that the bottom α-helix rather than the β3-β4 polybasic region in the C_2_B domain of Rph3A mediates SN25 interaction.

**Figure 4.**
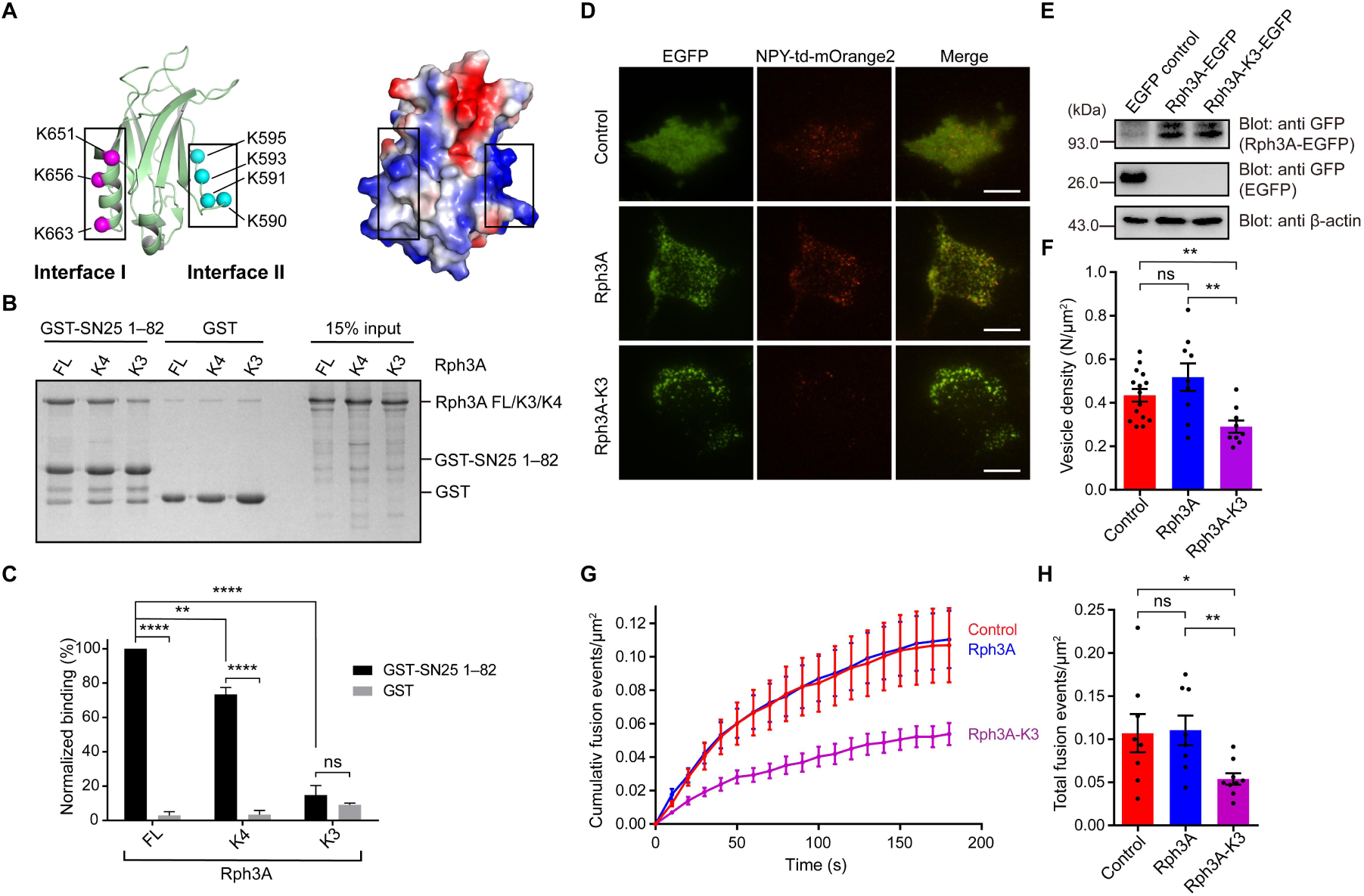
Importance of the C2B bottom α-helix of Rph3A FL for SN25 binding and DCV exocytosis in PC12 Cells. (A) Structural diagrams (left) and electrostatic surface potential (right) of Rph3A C2B (PDB entry: 5LOW). Interface I residues K651/K656/K663 on the bottom α-helix and interface II residues K590/K591/K593/K595 on the side are shown as magenta and cyan spheres, respectively. Black boxes display the basic patches that include the residues shown on the left. (B and C) Binding of Rph3A FL or mutations to the SN25 (1–82) measured by GST pull-down assay (B) and quantification of the Rph3A binding (C). K3, Rph3A FL protein bearing the K651A/K656A/K663A mutation in interface I; K4, Rph3A FL protein bearing the K590Q/K591Q/K593Q/K595Q mutation in interface II. Data are processed by Image J (NIH) and presented as the mean ± SEM (n = 3), technical replicates. (D) TIRF images of PC12 cells expressing EGFP control, Rph3A-EGFP or Rph3A-K3-EGFP mutant and DCV content (NPY-td-mOrange2). Scale bars, 10 μm. (E) Immunoblotting assay to determine the Rph3A, Rph3A-K3 mutant expression level in PC12 cells. Expression of EGFP control, Rph3A-EGFP, Rph3A-K3-EGFP and β-actin in PC12 cells was analyzed by 15% SDS-PAGE followed by immunoblotting with anti GFP antibody, and anti β-actin antibody, respectively. The positions of the molecular mass markers are shown on the left. (F) The density of docked vesicles was determined by counting the NPY-td-mOrange2 labeled vesicles in each image (n ≥ 9 cells in each). (G) NPY-td-mOrange2 release events detected by TIRF microscopy during sustaining high K+ stimulation. The curves indicate the cumulative number of fusion events per µm2 in each cell. (H) Quantification of the results at 180 s in the experiments of (G). Data are presented as mean ± SEM; (n ≥ 7 cells in each). Statistical significance and P values were determined by one-way (in F and H) or two-way (in C) analysis of variance (ANOVA). *P < 0.05; **P < 0.01; ****P < 0.0001; ns, not significant.

We next determined the functional importance of the bottom α-helix in DCV exocytosis in PC12 cells. Previous studies normally used exogenous overexpression approach to characterize the function of Rph3A. We then co-expressed Rph3A (or the Rph3A-K3 mutant) and NPY-td-mOrange2 in PC12 cells, and monitored the dynamics of NPY-td-mOrange2 labeled vesicles near the plasma membrane by TIRF microscopy as described in ***Figure 3***. Consistent with *in vitro* binding data, the numbers of plasma membrane-docked vesicles were significantly reduced in PC12 cells expressing the Rph3A-K3 mutant, compared to PC12 cells expressing Rph3A or control cells (***Figure 4D, F***). The distinct effects are unlikely due to the different protein expression levels in these transfected cells (***Figure 4E***). Similarly, the total number of release events in Rph3A-K3 expression cells was significantly fewer than Rph3A expression cells or control cells under high KCl stimulation (***Figure 4G, H***). Altogether, these results showed that the Rph3A–SN25 interaction mediated by the C_2_B bottom α-helix is essential for DCV docking and fusion of exocytosis in PC12 cells.

### Rph3A FL accelerates SNARE complex assembly via SN25 interaction

We next investigated how the Rph3A–SN25 interaction regulates DCV exocytosis. We explored the influence of the Rph3A–SN25 interaction on SNARE complex assembly by using FRET assay as previously described (Yang et al., 2015). Fluorescence-labeled Syb2 (29–93, S61C-BODIPY FL [BDPY], donor) and SN25 (1–206 or 11–206, R59C-Tetramethylrhodamine-5-maleimide [TMR], acceptor) were added with Syx1 (2–253), and SNARE complex assembly was detected by monitoring the decreased fluorescent signal of Syb2-BDPY due to FRET between fluorescence-labeled Syb2 and SN25 (***Figure 5A***). Intriguingly, we found that Rph3A FL remarkably accelerated SNARE complex assembly (***Figure 5B, C***). In contrast, the Rph3A-K4 mutant that maintains SN25 interaction accelerated the assembly as effectively as Rph3A FL, while the Rph3A-K3 mutant that impairs SN25 binding failed (***Figure 5B, C***). Hence, the assembly results suggest that the C_2_B bottom α-helix of Rph3A promotes SNARE complex assembly via SN25 binding.

**Figure 5.**
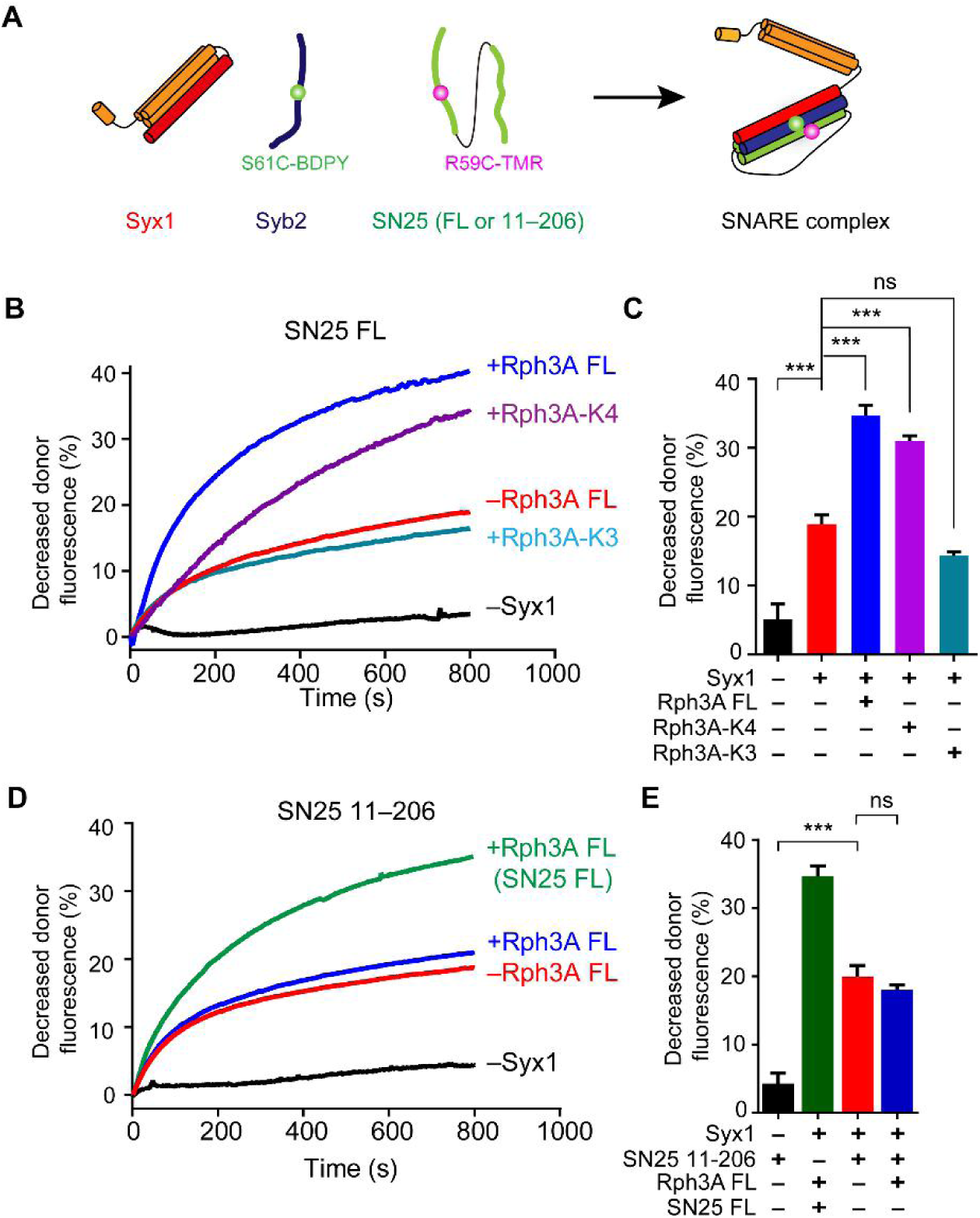
The function of Rph3A in SNARE complex assembly (A) Schematic diagram of SNARE complex assembly in the presence of Syx1 (2–253), Syb2 (29–93, S61C) and SN25 FL (1–206, R59C) or SN25 fragment (11–206, R59C). The FRET signal between Syb2 S61C-BDPY (donor) and SN25 R59C-TMR (acceptor) was monitored. (B) Effects of the K3 and/or K4 mutations on Rph3A-promoted SNARE complex assembly detected by FRET assay. (C) Quantification of the results at 600 s in the experiments of (B). (D) Effects on SN25 (11–206) on Rph3A-promoted SNARE complex assembly detected by FRET assay. Curve Rph3A (SN25 FL) (green color) represents the SN25 FL mediated SNARE complex assembly in the presence of Rph3A FL. (E) Quantification of the results at 600 s in the experiments of (D). Data are presented as mean ± SEM (n = 3), technical replicates. Statistical significance and P values were determined by one-way analysis of variance (ANOVA). ***P < 0.001; ns, not significant.

Since our data revealed that the N-peptide of SN25 mediates Rph3A FL interaction, we then explored the importance of the N-peptide of SN25 for the acceleration of SNARE complex assembly by Rph3A FL. Indeed, in contrast to SN25 FL, SN25 (11–206) that lacks the N-peptide strongly impaired Rph3A activity in accelerating the assembly (***Figure 5D, E***), demonstrating that Rph3A FL promotes SNARE complex assembly via binding of the C_2_B bottom α-helix to the N-peptide of SN25. Taken together, our data suggest that the Rph3A–SN25 interaction regulates DCV exocytosis via promoting SNARE complex assembly.

### Rph3A FL induces conformational change in the SN1 domain of SN25

Next, we explored the mechanism how the Rph3A–SN25 interaction promotes SNARE complex assembly. As the N-peptide of SN25 mediates Rph3A FL interaction, we suspect whether this interaction would induce potential conformational changes of the SNARE motif (SN1) adjacent to the N-peptide. To this aim, we used a bimane-tryptophan quenching assay to analyze conformational changes of SN1 of SN25, as this assay has been widely used to study the structure and movement of proteins and has shown strong sensitivity in short-distance electron transfer measurements (10 Å) (Mansoor et al., 2002; Islas and Zagotta, 2006; Taraska and Zagotta, 2010). In this case, a single tryptophan mutation (E55W) located at the middle portion of the SN1 was introduced in SN25 (full-length, 1–206), and a single cysteine mutation (R59C) adjacent to E55W was created for bimane labeling. If a conformational change occurred in SN1, for instance, a transition from random coils to α-helix, this change would shorten the distance between E55W and R59C-bimane thus producing enhanced bimane-tryptophan quenching (***Figure 6A***). In the original state of SN25 (unstructured state), SN25 E55W/R59C exhibited remarkably reduced bimane fluorescence compared to SN25 R59C (***Figure 6****–**figure supplement 1A***), indicating efficient electron transfer in the distance between E55W and R59C. As a positive control, addition of Syx1 and Syb2 to SN25 E55W/R59C led to further reduced bimane fluorescence (***Figure 6B, C***), consistent with the conformational change of SN1 (from random coils to α-helix) driven by the formation of the SNARE complex (Sutton et al., 1998). These data confirmed that conformational change of SN1 can be detected by using the bimane-tryptophan quenching assay. Intriguingly, when Rph3A FL was added to SN25 E55W/R59C in the absence of Syx1 and Syb2, we observed further reduced bimane fluorescence (***Figure 6B, C***), similar to that observed with the addition of Syx1 and Syb2. Hence, these results indicate that Rph3A is able to induce a conformational change of SN1 independent of Syx1 and Syb2.

**Figure 6.**
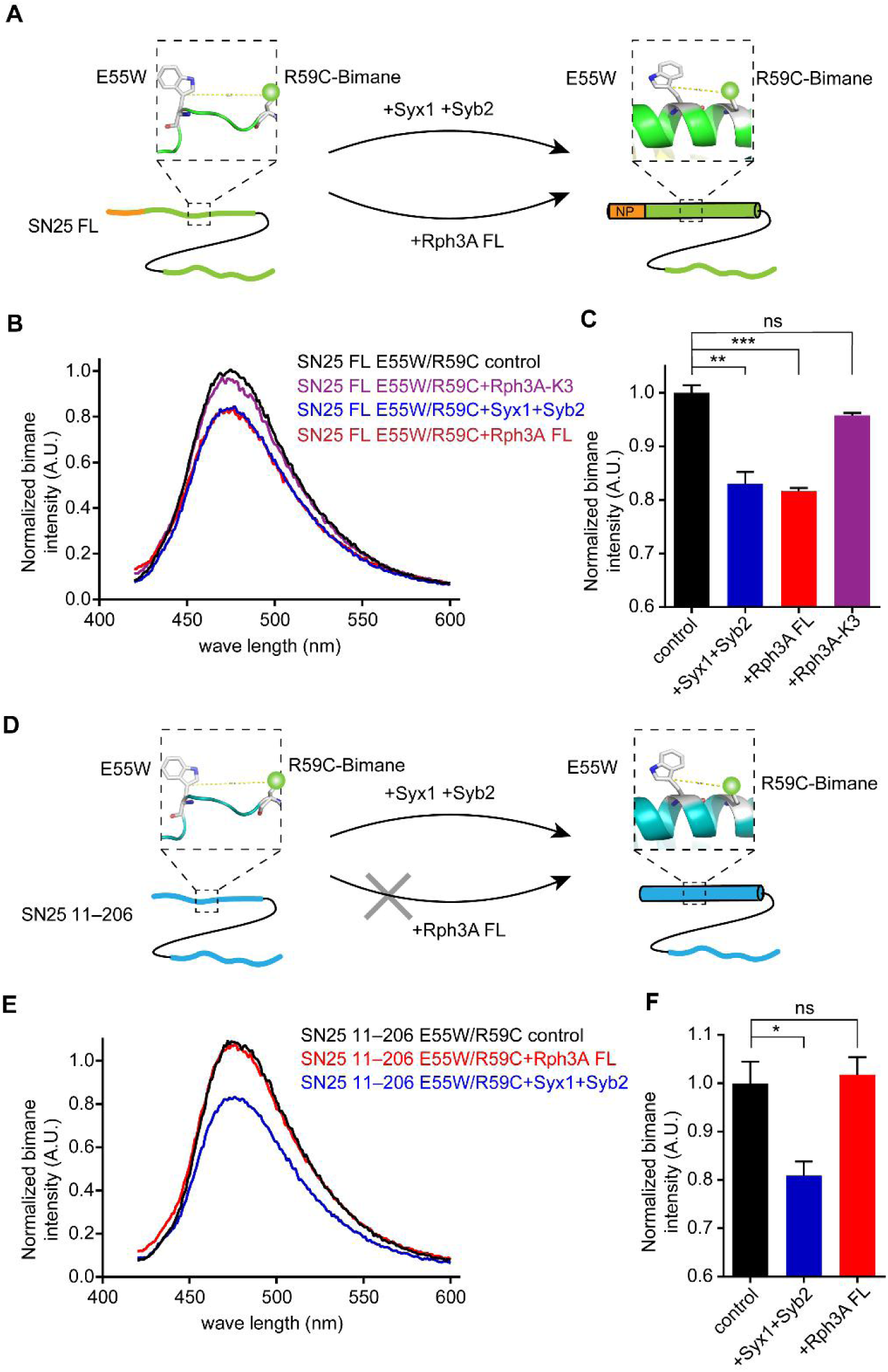
Conformation change of SN25 induced by Rph3A. (A and D) Illustration of the conformation change from random coils to α-helix in the SN1 of SN25 induced by Syx1 (2–253) and Syb2 (29–93), or by Rph3A, monitored by bimane-tryptophan quenching assay. Tryptophan was introduced at residue E55 and bimane fluorescence was labeled on residue R59C in SN25 FL (A) or SN25 11–206 (D). (B and C) Quenching of bimane fluorescence on the SN25 E55W/R59C with the addition of Syx1 and Syb2, and the addition of Rph3A FL or Rph3A-K3 mutant (B) and quantification of the results observed at 474 nm (C). (E and F) Quenching of bimane fluorescence on the SN25 (11–206) E55W/R59C with the addition of Syx1 and Syb2, or the addition of Rph3A FL (E) and quantification of the results observed at 474 nm (F). Data are presented as mean ± SEM (n = 3), technical replicates. The significant examined by one-way analysis of variance (ANOVA). *P < 0.05; **P < 0.01; ***P < 0.001; ns, not significant.

Then, we examined whether Rph3A FL-induced conformational change of SN1 is mediated by the interaction between the C_2_B bottom α-helix of Rph3A and the N-peptide of SN25. In this case, the Rph3A-K3 mutant and SN25 (11–206) were accordingly used in the bimane-tryptophan quenching assay. In contrast to Rph3A, addition of the Rph3A-K3 mutant to SN25 E55W/R59C failed to further reduce bimane fluorescence (***Figure 6B, C***), indicating the requirement of the C_2_B bottom α-helix of Rph3A. To verify the importance of the N-peptide of SN25, we created SN25 (11–206) R59C and SN25 (11–206) E55W/R59C (***Figure 6D***). As expected, SN25 (11–206) E55W/R59C exhibited reduced bimane fluorescence compared to SN25 (11–206) R59C (***Figure 6–figure supplement 1B***). Likewise, further reduced bimane fluorescence was observed for SN25 (11–206) E55W/R59C upon the addition of Syx1 and Syb2. However, in sharp contrast to SN25 E55W/R59C, SN25 (11–206) E55W/R59C displayed no further reduced bimane fluorescence upon the addition of Rph3A FL in the absence of Syx1 and Syb2 (***Figure 6E, F***), indicating the importance of the N-peptide of SN25. Taken together, our data suggest that Rph3A FL promotes SNARE complex assembly via inducing the conformational change of SN1 from random coil to α-helix, and the promotion requires the interaction of the C_2_B bottom α-helix of Rph3A with the N-peptide of SN25.

## Discussion

Rph3A was initially identified as a specific Rab3A/Rab27A-binding protein on secretory granules that is involved in the control of regulated vesicle secretion, including neurotransmitter release and hormone secretion (Li et al., 1994; Fukuda et al., 2004). Rph3A was later found to bind phospholipids via its C_2_ domains and interact with SN25 and phosphatidylinositol-4,5-bisphosphate (PI(4,5)P2) in via its C_2_B domain (Yamaguchi et al., 1993; Chung et al., 1998; Ferrer-Orta et al., 2017). In particular, the C_2_B domain of Rph3A was generally regarded as the pivotal unit to bind SN25, and this interaction was implicated to regulate docking and fusion of DCVs in neuroendocrine cells and to control repriming of synaptic vesicles in neurons (Tsuboi and Fukuda, 2005; Deak et al., 2006). Despite these advances, the physiologically relevant binds sites between Rph3A and SN25 remain elusive, and how the Rph3A–SN25 interaction regulates exocytosis has not been fully elucidated. In the present study, we found a new binding mode between Rph3A and SN25, and revealed its importance in regulating DCV exocytosis in PC12 cells. Our data highlight that this binding mode is essential for Rph3A activity in promoting SNARE complex assembly via a mechanism involving the conformational change of SN25.

The Rph3A C_2_B domain was previously identified to bind SN25 (Tsuboi and Fukuda, 2005; Deak et al., 2006), but the physiologically relevant binding sites remain elusive. Our present data for the first time identified an Rph3A–SN25 interaction that relies on the intramolecular interplay between the RBD and C_2_AB of Rph3A (***Figure 1***), suggesting that both domains act synergistically to bind SN25. In addition, this interaction is compatible with Rab3A interaction, indicative of the formation of a Rab3A–Rph3A–SN25 ternary complex essential for bridging vesicles and the plasma membrane (***Figure 7***). Besides, considering that the PRL of Rph3A connecting the RBD and C_2_AB domains contains multiple sites which can be phosphorylated by PKA, PKC and/or CaMKII (Kato et al., 1994; Numata et al., 1994; Fykse et al., 1995; Foletti et al., 2001), it is plausible that phosphorylation of the PRL is able to alter the Rph3A–SN25 interaction via influencing the intramolecular interplay of the RBD with C_2_AB, which modulating Rph3A activity *in vivo*.

**Figure 7.**
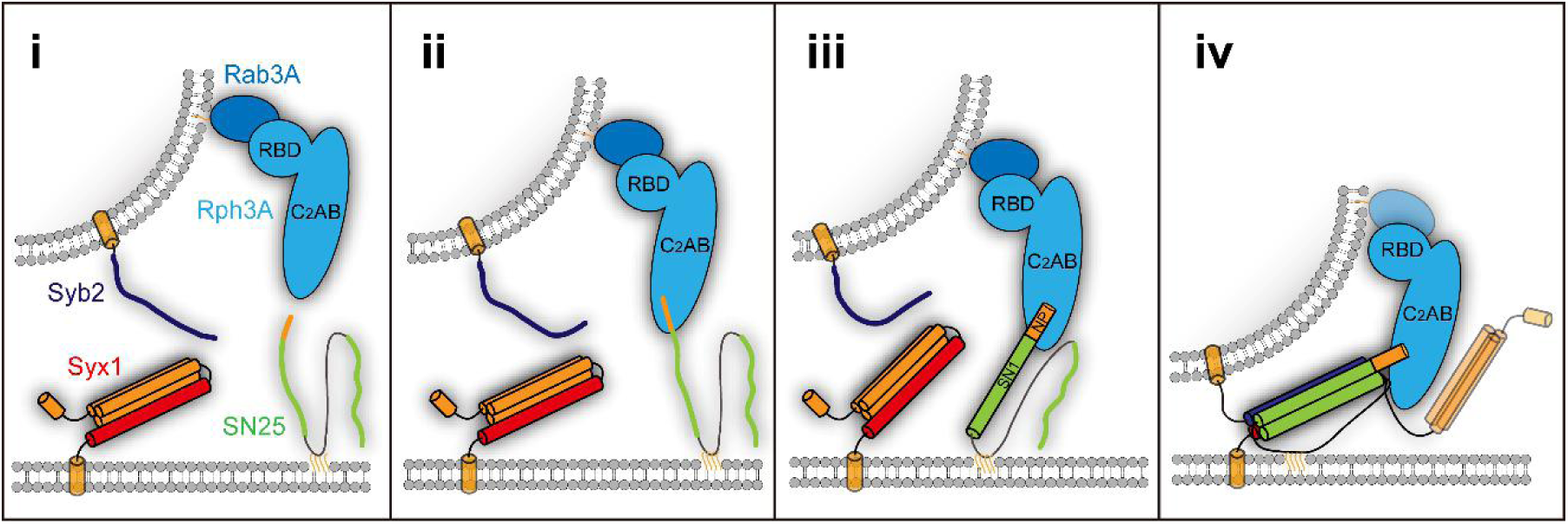
Model of Rph3A function in prefusion steps of exocytosis. Stage i, Rph3A is associated with trafficking vesicles via binding to GTP-Rab3A. Stage ii, Rph3A promotes vesicle docking via binding to plasma membrane-bound SN25 and vesicle-bound Rab3A together. Stage iii, upon binding to the N-peptide of SN25, Rph3A induces a conformational change of SN1 from random coils to α-helix. Stage iv, Rph3A accelerates SNARE complex assembly via SN25 interaction, which promote vesicle priming. As indicated, multiple SNARE regulatory proteins, e.g., Munc18-1, Munc13-1, Doc2, and Syt1 are also involved in the stage i–iv.

In addition to the SNARE motifs of SN25 involved in the formation of the SNARE four helical bundle, other sequences in SN25 have gained attention in recent years. For instance, the linker that connects SN1 and SN2 has been suggested to act as a flexible molecular spacer to ensure efficient S-acylation and support fusion intermediates by local lipid interactions (Salaun et al., 2020; Shaaban et al., 2019), or recruited by Munc13-1 to chaperone SNARE complex assembly (Kalyana Sundaram et al., 2021). In the present study, we identified that the N-peptide sequence (1–10) rather than the SNARE motifs of SN25 mediates interaction with Rph3A FL (***Figure 2***), and this binding specificity might be conferred by synergistic action of the RBD with the C_2_AB of Rph3A as discussed above.

Actually, according to the recently solved Rph3A-C_2_B/SN25 (Δ1–6) structure (Ferrer-Orta et al., 2017), three distinct regions in SN25 were shown to participate in binding to the bottom α-helix of Rph3A C_2_B domain, one involved R8/E10/Q15 in the N-terminal peptide of SN25, the other two included E38/D41/R45 and D51/E55/R59 in the middle of SN1, respectively. However, these binding information need to be interpreted with caution, because in previous work i) the isolated C_2_B domain, but not Rph3A FL was employed; ii) the N-peptide residues of SN25 were partially deleted; and iii) physiological relevance of these interactions had not been tested yet. Our observations that Rph3A FL bound efficiently to SN25 E38A/D41A/R45A mutant and D51A/E55A/R59A mutant but not to SN25 lacking the N-peptide pinpoint the binding specificity between Rph3A FL and the SN25 N-peptide (***Figure 2E*** *and* ***Figure 2–figure supplement 1***). Consistently, deletion of the N-peptide of SN25 or disruption of the C_2_B bottom α-helix of Rph3A significantly impaired docking and fusion of DCVs with the plasma membrane in PC12 cells (***Figure 3*** *and* ***4***), revealing the physiological relevance of such binding mode.

Doc2 and Synaptotagmin-1(Syt1), which contain tandem C_2_ domains homologous to that of Rph3A, are major Ca^2+^ sensors in neurons responsible for spontaneous, asynchronous and synchronous release of neurotransmitters (Kaeser and Regehr, 2014). Doc2 and Syt1 were both implicated to bind SN25 at the middle of SN1 (D51/E52/E55) (Groffen et al., 2010; Brewer et al., 2015; Zhou et al., 2015). In this circumstance, the preference for binding of Rph3A to the N-peptide of SN25 identified here could make D51/E52/E55 more available for Doc2 and/or Syt1 interaction. In addition, interestingly, our results found for the first time that Rph3A has an ability to promote SNARE complex assembly, which requires binding of the C_2_B bottom α-helix of Rph3A to the N-peptide of SN25 (***Figure 5***). As indicated by bimane-tryptophan quenching assay (***Figure 6***), our results found that Rph3A induces a conformation change in the SNARE motif of SN25, which increases the content of SN1’ α-helical configuration to facilitate its assembly with the SNARE motifs of Syx1 and Syb2. Taken together, we suggest that Rph3A has at least two important roles in prefusion events of exocytosis: one is to promote vesicle docking dependent on its associations with Rab3A and SN25 between the two opposite membranes; the other is to promote vesicle priming via accelerating SNARE complex assembly (***Figure 7***). These mechanisms may partly underlie the functional importance of Rph3A in DCV exocytosis in PC12 cells.

Previous studies have revealed that the N-peptide of Syx1 bound to Munc18-1 plays a regulatory role in exocytosis and that the N-terminal proline-rich sequence of Syb2 bound to intersectin-1 is necessary for the clearance of SNARE complexes required for vesicle recycling (Rathore et al., 2010; Japel et al., 2020). Combined with these results, the functional importance of the N-peptide of SN25 bound to Rph3A identified here suggests a common feature that the N-terminal regions of the three SNAREs are required for modulating vesicle exocytosis via interactions with multiple regulatory proteins.

In isolated form, the SNARE motifs of Syx1, SN25 and Syb2 all assume unstructured conformation (Fasshauer et al., 1997; Hazzard et al., 1999). However, as observed in the crystal structures of Munc18-1 bound to Syb2 or Syx1 and of Vps33 bound to Nyv1 or Vam3, the SNARE motifs of Syb2 or Syx1, and Nyv1 or Vam3 display highly ordered α-helical structures (Baker et al., 2015; Stepien et al., 2022), resemblance to their conformations in the assembled SNARE complex, suggesting that the prestructured SNARE motif of the SNAREs might be essential for proper SNARE pairing and register. Highly consistent with this notion, our data showed that Rph3A bound to SN25 renders SN1 to assume a prestructured conformation (Fig. 6). These evidence suggests that the prestructured state of the three SNAREs induced by multiple regulatory proteins might be a prerequisite for the efficiency of SNARE complex assembly in exocytosis.

## Materials and Methods

### Plasmids and protein purification

The rat Syb2 (29–93) and Syb2 (29–93, S61C), mouse Rab3A (22–217, Q81L) were all cloned into pGEX-6p-1(GE Healthcare; Piscataway, NJ). The cytoplasmic domain of rat Syx1 (2–253) was cloned into pGEX-KG vector. These proteins were expressed and purified as described (Dulubova et al., 2005; Ma et al., 2013; Stepien et al., 2022).

The full-length human SN25 (1–206), a series of fragments 1–82, 1–140, 141–206, 83–206, 83–140, N-peptide deletion mutants 7–82 and 11–82, the SN25 1–82 EDR mutant (E38A/D41A/R45A) and DER mutant (D51A/E55A/R59A), the rat Rph3A RBD (40–170) and RBD G102A/L104A mutant were all cloned into pGEX-6P-1 vector, incorporating an N-terminal PreScission protease-cleavable GST tag. The human SN25 (1–206, R59C), SN25 (1–206, E55W/R59C), SN25 (11–206, R59C) and SN25 (11–206, E55W/R59C) were cloned into pET28a vector (Novagen, Australia), incorporating an N-terminal His_6_ tag. All the recombinant proteins were expressed in *E. coli* BL21 (DE3) strain cultured in LB media at 37°C to OD600 of 0.6–0.8 and were induced with 0.4 mM IPTG at 20°C for 16 hrs. Cells were resuspended in a buffer containing 25 mM HEPES pH 7.4, 150 mM KCl, 10% glycerol (v/v). Cells were broken using AH-1500 Nano Homogenize Machine (ATS Engineering Inc.) at 800 bars three times at 4°C. Cell lysates were centrifuged at 16,000 rpm for 30 minutes in a JA-25.50 rotor (Beckman Coulter) at 4°C. For the purification of GST fusion proteins, the supernatants were incubated with 2 mL glutathione-Sepharose beads (GE Healthcare) at 4°C for 3 hrs. The bound proteins were eluted by a buffer containing 20 mM Tris pH 8.0, 150mM NaCl and 20mM L-glutathione at 4°C for 3 hrs. For the purification of His_6_ fusion proteins, the supernatant was incubated with Ni^2+^-NTA agarose (QIAGEN) at 4°C for 1 hr. The beads were washed with buffer 25 mM HEPES pH 7.4, 150 mM KCl, 10% glycerol (v/v) supplied with an additional 30 mM imidazole. The protein was eluted with a wash buffer described above but supplied with an additional 300 mM imidazole.

The full-length Rph3A (1–681) gene was amplified from rat brain cDNA library and cloned into the pEXP5-NT/TOPO vector (Invitrogen). The Rph3A fragments 1–281, 182–681, 282–681, and 372–681 were cloned into the pET28a vector. The Rph3A middle PRL loop deletion fragments and mutants Rph3A (Δ185–371), Rph3A (Δ185–371) G102A/L104A, Rph3A (Δ1–45 &Δ161–371), Rph3A-K3 mutant (K651A/K656A/K663A) and Rph3A-K4 mutant (K590Q/K591Q/K593Q/K595Q) were also cloned into the pEXP5-NT/TOPO vector. For expression of full-length Rph3A and its truncation or mutants via pEXP5-NT/TOPO or pET28a vector, plasmids were transformed into *E. coli* BL21 (DE3). Cells were grown to OD 600 of 0.6 in 37°C and IPTG was added to a final concentration of 0.2 mM. Cells were shaken overnight at 16°C for 18 hrs, pelleted and resuspended in lysis buffer (25 mM HEPES pH 7.4, 1M KCl). Cells were lysed by AH-1500 Nano Homogenize Machine (ATS Engineering Inc.). Triton X-100 was added to a final concentration of 0.2% and PMSF to 1 mM. Lysates were incubated with 1 ml of Ni-NTA resin (QIAGEN) in 4°C for 1 hr. The resin was washed with 0.1 M citrate buffer (3 mM citric acid, 97 mM trisodium citrate, pH 6.3) containing 30 mM imidazole, and the bound Rph3A proteins were eluted with 300 mM imidazole in citrate buffer. All the eluted proteins were loaded into Superdex 200 pg or Superdex 75 pg size exclusion chromatography (GE Healthcare) to remove aggregates and potential contaminants.

The plasmid encoding NPY-td-mOrange2 was purchased from Addgene (plasmid no. 83497). For the SN25 knockdown in PC12 cells, the SN25 short hairpin RNA (shRNA) was designed as reported (Cahill et al., 2006) and cloned into pFHUUIG_shortU6 (L309) plasmid after the H1 promoter. The SN25 FL (1–206) or truncation fragment (11–206) rescue genes were cloned into pFHUUIG_shortU6 (L309) plasmid after the Ub promoter. Full-length rat Rph3A and Rph3A-K3 mutation were cloned into pEGFP-N3 vector in ***Figure 4***.

### Analytical ultracentrifugation (AUC)

To identify the aggregation state of Rph3A, sedimentation velocity (SV) experiments were performed using ProteomeLab XL-I analytical ultracentrifuge (Beckman Coulter, Palo Alto, CA, USA). Rph3A proteins were prepared in 0.1 M Citrate buffer. The high purity proteins were collected and then concentrated to 1 OD (UV, 280 nm). Before the sample loading, a highly speed centrifuge (12,000 rpm) was done to remove the denatured protein aggregations. The reference cell was loaded with 400 μl 0.1 M Citrate buffer, and the sample cell was loaded with 380 μL Rph3A protein samples. Experiments were done with 40,000 rpm speed at 4°C overnight (An-60 Ti Rotor). Sedimentation profiles were recorded with UV (280 nm) absorbance and interference optics (660 nm), and these data were analyzed by SEDFIT software (Schuck, 2000) and plotted with Prism 8.0.0.

### GST pull**-**down assay

Firstly, to reduce the non-specific protein binding, the Glutathione Sepharose 4B affinity beads (GST beads) (GE Healthcare) were washed three times with buffer 25 mM HEPES pH 7.4, 150m M KCl containing 0.1% bovine serum albumin (BSA) and incubated at 4°C overnight. Then, 4 μM GST fused proteins (GST-SN25 FL or mutants) were incubated with 5 μM different Rph3A fragments or other proteins and 20 μl 50% (v/v) GST beads to a final volume of 100 μl with buffer 25 mM HEPES pH 7.4, 150 mM KCl containing 0.1% BSA. 4 μM purified GST-free proteins were used in the control groups. After gentle shaking at 4°C for 3 hrs, beads were washed three times using buffer 25 mM HEPES pH 7.4, 150 mM KCl with 500 ×g, 10 min. The bound proteins were eluted with 32 μl buffer 20 mM Tris pH 8.0, 150 mM NaCl containing 20 mM L-glutathione. Samples were separated by SDS-PAGE and the protein bands were detected by Coomassie blue staining. In ***Figure 1G***, the bound proteins were analyzed by Western-Blot and probed with mouse monoclonal anti-His_6_ (Proteintech; 66005-1-lg). Each experiment was repeated at least three times. Data were analyzed by ImageJ and Prism 8.0.0.

### DCVs docking and secretion assay in PC12 Cells

For SN25 knockdown in PC12 cells, the lentiviral expression vectors (control L309 plasmids, L309-SN25 shRNA plasmids, L309-SN25 FL or L309-SN25 11–206 rescue plasmids) were individually co-transfected with three helper plasmids (pRSVREV, pMDLg/pRRE, and pVSVG) into HEK293T cells as described before (Wang et al., 2020). The lentiviruses were harvested after 48 hrs and concentrated by sucrose density gradient centrifugation, finally resuspended with 20 μl PBS buffer.

For the DCV docking and secretion assay, PC12 cells were cultured in RPMI Medium 1640 basic (1×) (GIBCO) supplemented with 10% FBS (GIBCO) at 37°C in a 5% CO2 atmosphere at constant humidity. For TIRF imaging in ***Figure 3***, PC12 cells were plated on 20 mm glass bottom dish (NEST, China). Then, 1 μg of NPY-td-mOrange2 plasmids were co-transfected with 20 μl prepared SN25 knockdown or rescue lentiviruses by using LipofectAmine 3000 (Invitrogen, USA) according to the manufacturer’s instructions. After 48–72 hrs, cells were imaged by TIRF microscopy on a Nikon Ti inverted microscope equipped with a 100× oil-immerse objective (NA 1.49) and an EMCCD camera. To analyze the DCV docking, the imaging was performed in a basal buffer (15 mM HEPES, pH 7.4, 145 mM NaCl, 5.6 mM KCl, 2.2 mM CaCl2, 0.5 mM MgCl2, 5.6 mM glucose), and the docked DCVs were monitored by NPY-td-mOrange2. Spot detector plugin of Icy software were used to analyze the number of docked DCVs. In order to monitor the number of fusion events in per cell, a common highly K^+^ stimulation buffer (15 mM HEPES, pH 7.4, 95 mM NaCl, 60 mM KCl, 2.2 mM CaCl2, 0.5 mM MgCl2, 5.6 mM glucose) was used to displace the basal buffer to depolarize PC12 cells and trigger vesicles fusion. We monitored exocytosis of NPY-td-mOrange2 at the single-vesicle level as described before (Zhou et al., 2019). Exocytosis events were scored manually. Orange fluorescence was excited with a 532 nm laser, and EGFP fluorescence was excited with a 488 nm laser. Images were analyzed by NIS-Elements viewer 4.20 and Prism 8.0.0 software.

In ***Figure 4***, PC12 cells were co-transfected 1 μg of NPY-td-mOrange2 and 1 μg of pEGFP-N3, pEGFP-N3-Rph3A, pEGFP-N3-Rph3A-K3 by using LipofectAmine 3000 (Invitrogen, USA) according to the manufacturer’s instructions. The DCV docking and secretion assay were performed as above.

### Immunoblotting

The cell culture procedure was the same as above. In ***Figure 3A*** and ***Figure 4E***, cells were harvested at 72 h after transfection. Then, collected cells were solubilized with 100 μl RIPA lysis buffer in 4°C for 1 hr. The insoluble components were removed by 12,000×g, 10 min and the supernatants were boiled with 5× SDS loading buffer for 10 min. Equal quantities of protein were subjected to 15% SDS-PAGE followed by immunoblotting with anti-SN25 antibody (Proteintech; 14903-1-AP) (***Figure 3A***), anti GFP antibody (Proteintech; 66002-1-Ig) (***Figure 4E***), and anti β-actin antibody (Proteintech; 66009-1-Ig). Immunoreactive bands were visualized either with HRP-conjugated goat anti-mouse (Proteintech; SA00001-1) or with anti-rabbit IgG (Proteintech; SA00001-2) and detected by ECL reagent.

### Fluorescence resonance energy transfer (FRET) assay

For the SNARE complex assembly assay in ***Figure 5***, the protein labeling procedure was performed as described before (Yang et al., 2015). Purified Syb2 (29–96, S61C) was labeled with 6× molar excess BODIPY FL N-(2-aminoethyl)-maleimide (BDPY) (Molecular Probes) and SN25 FL (1–206, R59C) or truncation mutant (11–206, R59C) were labeled with 5×molar excess Tetramethylrhodamine-5-maleimide (TMR) (Molecular Probes) in 25 mM HEPES pH 7.4, 150 mM KCl buffer in 4°C for overnight. The labeling reaction was stopped by 10 mM DTT. Excess fluorescent dyes were removed by a PD-10 desalting column (GE Healthcare) in buffer 25 mM HEPES pH 7.4, 150 mM KCl. In ***Figure 5B, D***, 2 μM Syx1 (2–253) and 1 μM Syb2 S61C-BDPY were incorporated into a mixture of TMR labeled SN25 FL or 11–206 (2 μM) and Rph3A WT or mutants (8 μM). FRET assays were carried out with a PTI QM40 fluorescence spectrophotometer at 20°C using a 1-cm quartz cuvette. Donor fluorescence was monitored with excitation and emission wavelength of 485 nm and 513 nm, respectively. Each experiment was repeated at least three times. The decreased donor fluorescence intensity was analyzed by Prism 8.0.0 software.

### Bimane-tryptophan quenching assay

Purified SN25 FL or 11–206 (E55W/R59C) was labeled with 10× molar excess monobrombimane (mBBr, Molecular Probes; Eugene, OR) in buffer 25 mM HEPES pH 7.4, 150 mM KCl containing 0.2 mM TCEP, 0.2 mM EDTA. After incubation at 4°C overnight, reactions were stopped by adding 10 mM DTT. Excess fluorescent dyes were removed by a PD-10 desalting column (GE Healthcare) with buffer 25 mM HEPES pH 7.4, 150 mM KCl. In this assay, 0.7 μM bimane-labeled SN25 FL or 11–206 (E55W/R59C) were mixed with 5.6 μM Rph3A or 1.4 μM Syx1 (2–253) and 1.4 μM Syb2 (29–93). Experiments were performed at 20°C in buffer A (25 mM HEPES pH 7.4, 150 mM KCl). The bimane fluorescence was excited with a 380 nm wavelength, and monitored with an emission wave scan from 400 nm to 600 nm on a PTI QM-40 spectrofluorometer. SN25 FL or 11–206 R59C mutation used in ***Figure 6–figure supplement 1*** were also labeled bimane fluorescence as above. Each experiment was repeated at least three times.

#### Sequence alignment

The sequence alignment was performed using Clustal Omega.

#### Statistical analysis

Prism 8.0.0 (GraphPad) and Image J (NIH) were used for graphing and statistical tests.

## Data availability

All data generated or analyzed during this study are included in the manuscript and supporting file; Source data files have been provided for Figures 1-6 and Figure supplements.

## Acknowledgments

We thank Josep Rizo (University of Texas Southwestern Medical Center at Dallas) for providing Rab3A Q81L construct. We thank Andreas Mayer (Department of Biochemistry, University of Lausanne) for providing pEXP5-NT/TOPO vector. We thank Zhengxing Wu (Huazhong University of Science and Technology) for sharing the EMCCD camera.

## Supplementary figures

**Figure 1-figure supplement 1.**
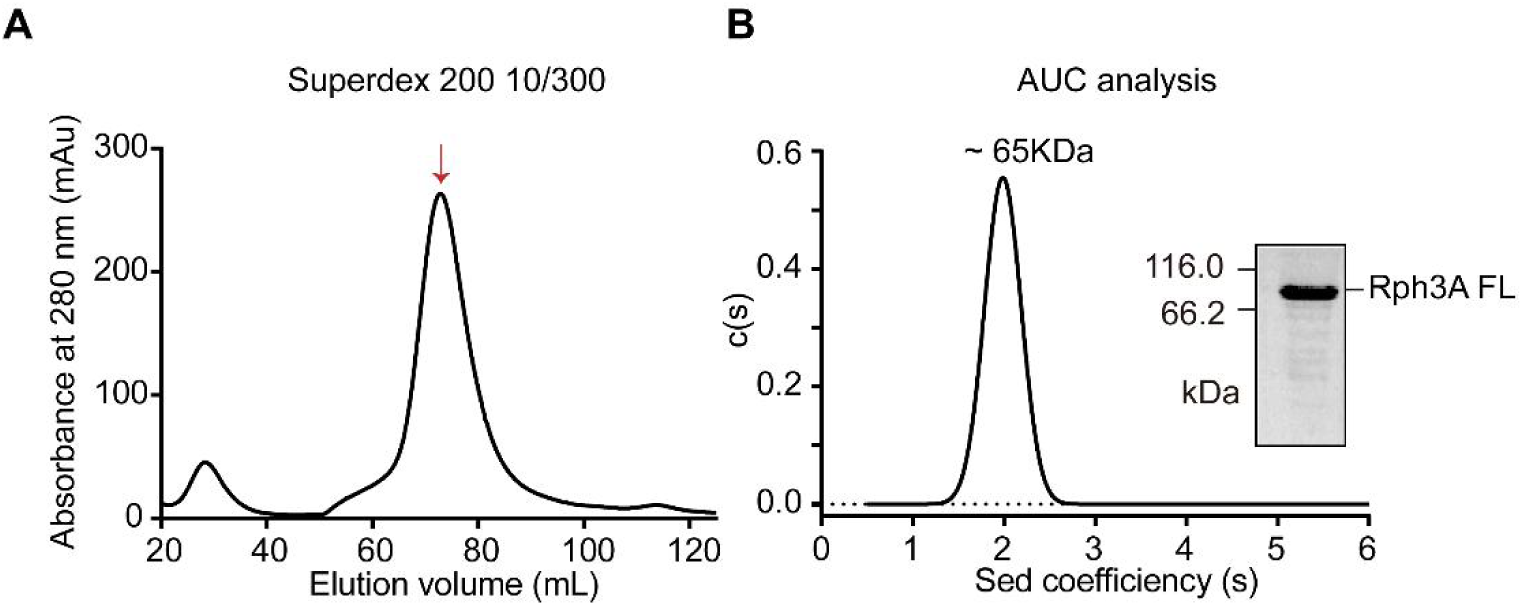
Size-exclusion chromatograph and analytical ultracentrifugation analysis of Rph3A FL protein. (A) Size-exclusion chromatograph (SEC) elution profiles of Rph3A FL protein with Superdex 200 10/300 GL, red arrow represents the elution peak of Rph3A FL protein. (B) SV analytical ultracentrifugation (AUC) analysis of Rph3A FL protein (left), the molecular weights were calculated based on sedimentation coefficient, obtained with SEDFIT software with a continuous c(s) model. The right panel represents Rph3A FL sample before the SV assay, resolved with SDS-PAGE.

**Figure 1-figure supplement 2.**
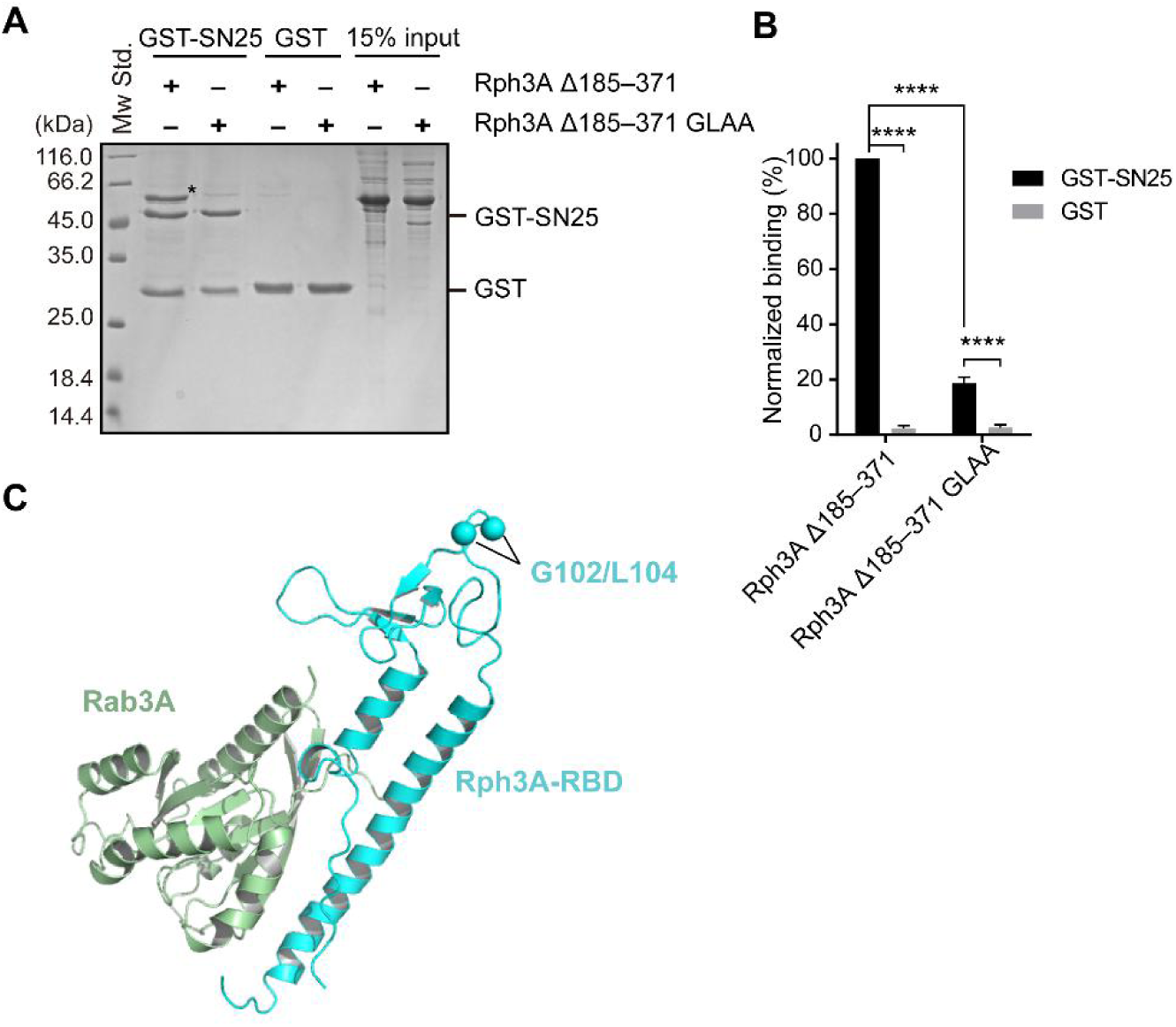
The effect of G102A/L104A mutant on Rph3A–SN25 interaction. (A) Binding of Rph3A Δ185–371 WT or G102A/L104A mutant to GST-SN25 measured by GST pull-down assay. Asterisk shows a band of bound protein. (B) Quantification of bound Rph3A Δ185–371 proteins in (A). Data are processed by Image J (NIH) and presented as the mean ± SEM (n = 3), technical replicates. Statistical significance and P values were determined by one-way analysis of variance (ANOVA). ****P < 0.0001. (C). The structure of Rph3A-RBD–Rab3A complex (PDB entry: 1ZBD). The G102/L104 site was mapped to Rph3A-RBD domain.

**Figure 2–figure supplement 1.**
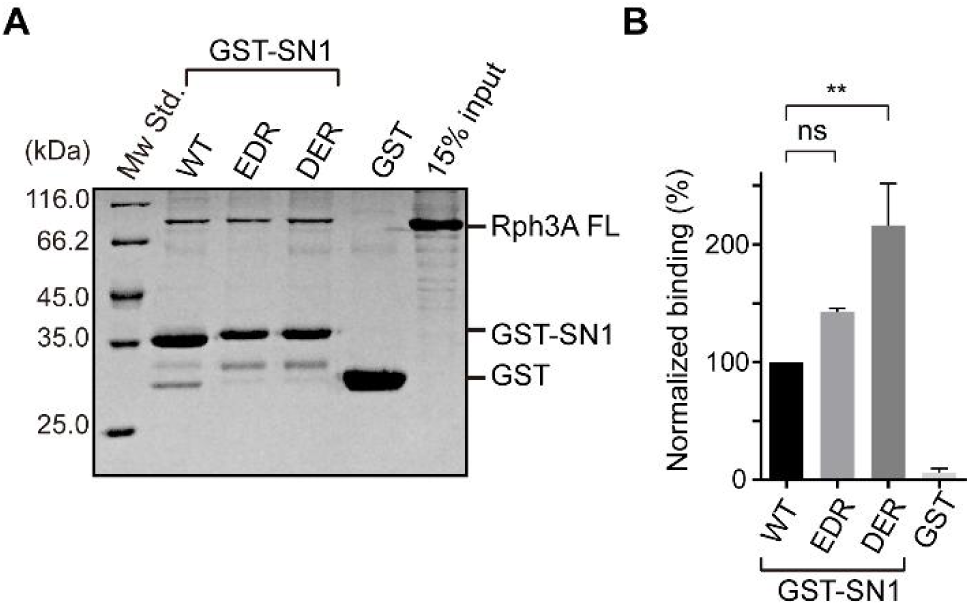
Effect of middle region mutants in SN1 on Rph3A interaction. (A) GST pull-down assays were performed with GST-SN1 WT or EDR (E38A/D41A/R45A), DER (D51A/E55A/R59A) mutants and Rph3A FL protein. (B) Quantification of the relatively bound Rph3A shown in (A). All data is presented as mean ± SEM (n = 3), technical replicates. Statistical significance and P values were determined by one-way analysis of variance (ANOVA). **P < 0.01; ns: not significant.

**Figure 2–figure supplement 2.**
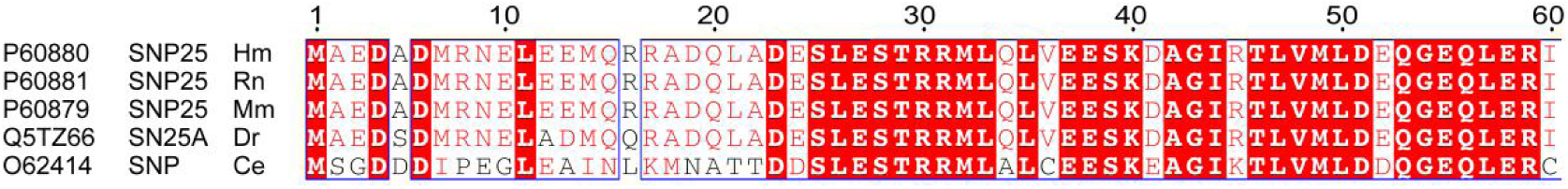
Alignment of the N-terminus sequences of SN25 across species using Clustal Omega Program and Escript 3.0. Uniprot entries are shown on the left of the chart. Residues with 100% consensus are shaded by red, residues which consensus level > 80% are colored red. *Hs*, *Homo sapiens*; *Rn*, *Rattus norvegicus*; *Mm*, *Mus musculus*; *Dr*, *Danio rerio*; *Ce*, *Caenorhabditis elegans*.

**Figure 4–figure supplement 1.**
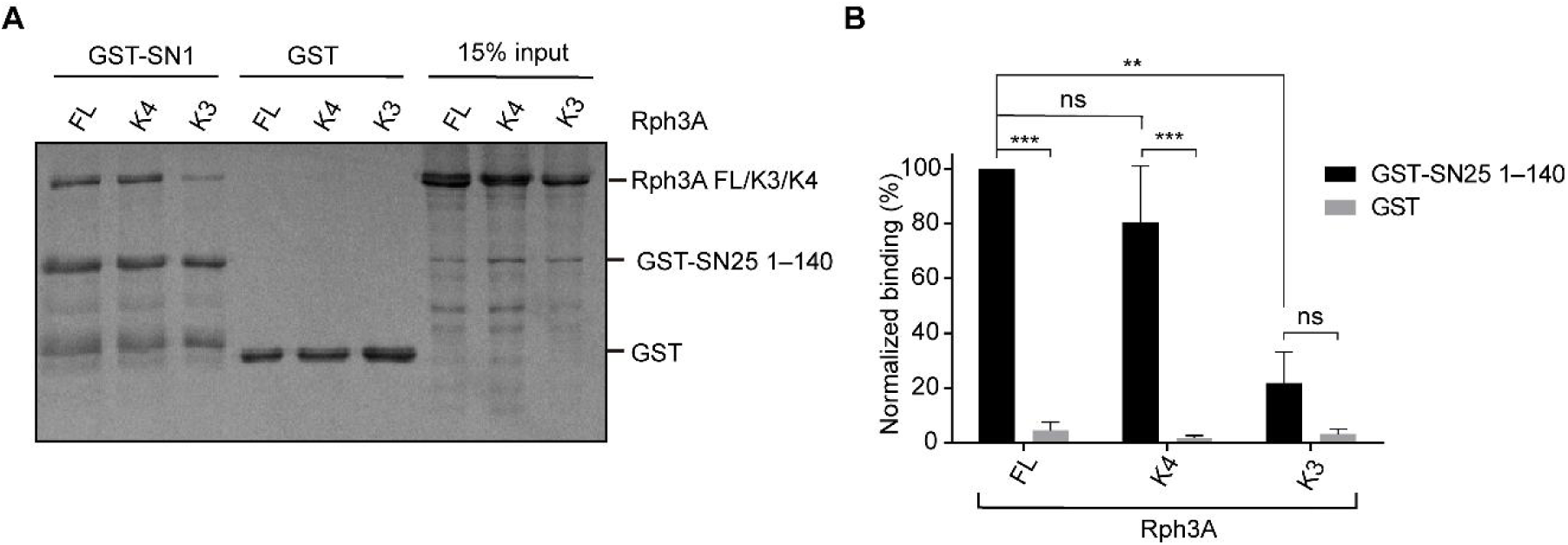
Effect of Rph3A interface I mutant on SN25 (1–140) binding. (*A* and *B*) Binding of Rph3A WT or K3, K4 mutants to the GST-SN25 (1–140) measured by GST pull-down assay (*A*) and quantification of the Rph3A binding (*B*). All data is presented as mean ± SEM (n = 3), technical replicates. Statistical significance and P values were determined by one-way analysis of variance (ANOVA). \*\**P* < 0.01; ns: not significant.

**Figure 6–figure supplement 1.**
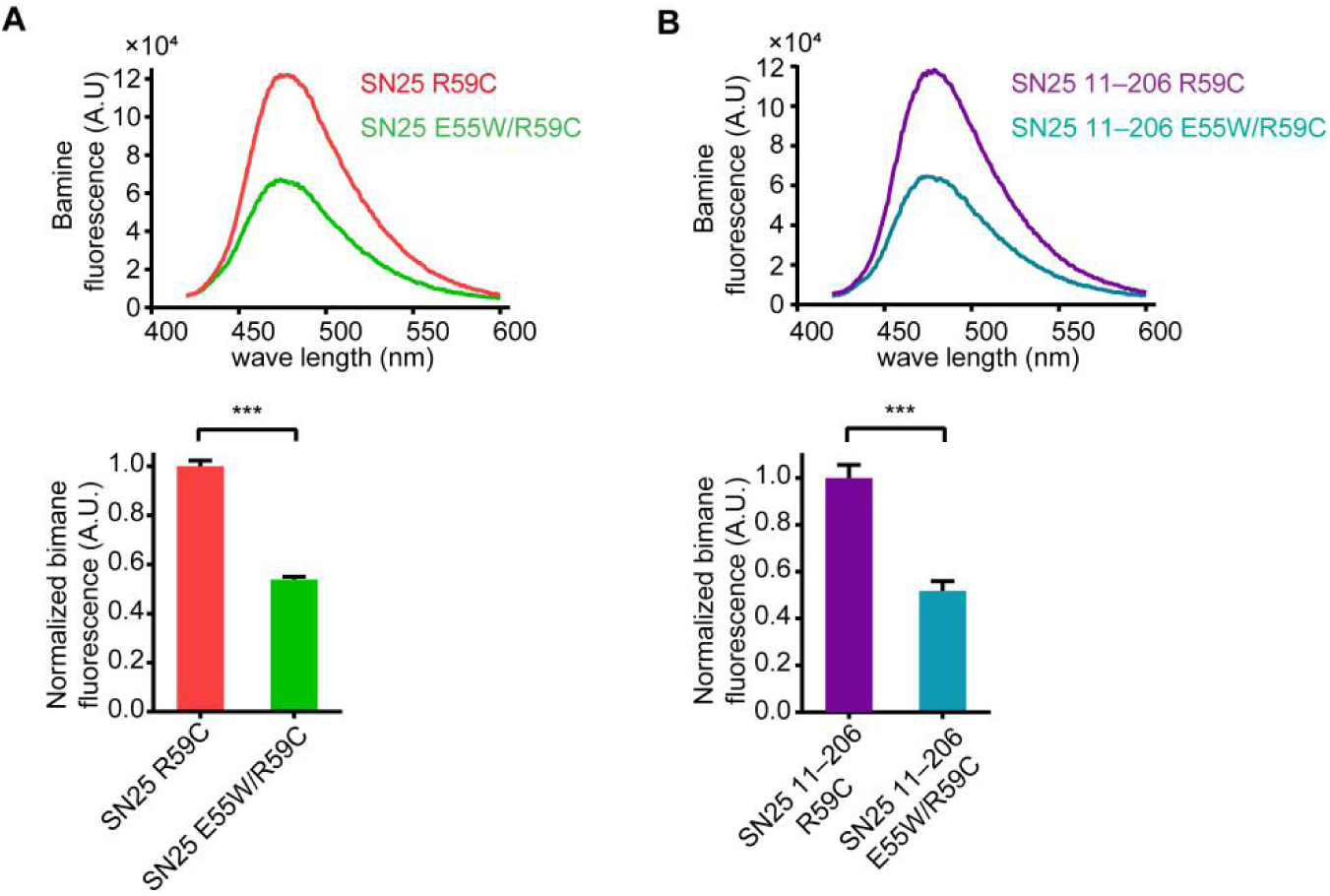
The intrinsic bimane fluorescence quenching by E55W mutant on SN25. (*A*) Comparison of bimane fluorescence between SN25 R59C labeling and SN25 E55W/R59C labeling. (*B*) Comparison of bimane fluorescence between SN25 11–206 R59C labeling and SN25 11–206 E55W/R59C labeling. Quantification of the bimane fluorescence observed at 474 nm in (*A* and *B*) is shown on the bottom. All data is presented as mean ± SEM (n = 3), technical replicates. Differences among groups by unpaired Student’s t-test. \*\*\**P* < 0.001.

## Source data

Figure 1-source data 1: Uncropped SDS-gel shown in figure 1B.

Figure 1-source data 2: Excel file with data used to make figure 1C, 1E, 1H and 1J.

Figure 1-source data 3: Uncropped SDS-gel shown in figure 1D.

Figure 1-source data 4: Uncropped SDS-gel and Western blot shown in figure 1G.

Figure 1-source data 5: Uncropped SDS-gel shown in figure 1I.

Figure 2-source data 1: Uncropped SDS-gel shown in figure 2B.

Figure 2-source data 2: Excel file with data used to make figure 2C and 2F.

Figure 2-source data 3: Uncropped SDS-gel shown in figure 2E.

Figure 3-source data 1: Uncropped Western blot shown in figure 3A.

Figure 3-source data 2: Excel file with data used to make figure 3C-E.

Figure 4-source data 1: Uncropped SDS-gel shown in figure 4B.

Figure 4-source data 2: Excel file with data used to make figure 4C and 4F-H.

Figure 4-source data 3: Uncropped Western blot shown in figure 4E.

Figure 5-source data 1: Excel file with data used to make figure 5C and 5E.

Figure 6-source data 1: Excel file with data used to make figure 6C and 6F.

Figure 1-figure supplement 1-source data 1: Uncropped SDS-gel shown in Figure 1-figure supplement 1B.

Figure 1-figure supplement 2-source data 1: Uncropped SDS-gel shown in Figure 1-figure supplement 2A.

Figure 1-figure supplement 2-source data 2: Excel file with data used to make Figure 1-figure supplement 2B.

Figure 2-figure supplement 1-source data 1: Uncropped SDS-gel shown in Figure 2-figure supplement 1A.

Figure 2-figure supplement 1-source data 2: Excel file with data used to make Figure 2-figure supplement 1B.

Figure 4-figure supplement 1-source data 1: Uncropped SDS-gel shown in Figure 4-figure supplement 1A.

Figure 4-figure supplement 1-source data 2: Excel file with data used to make Figure 4-figure supplement 1B.

Figure 6-figure supplement 1-source data 1: Excel file with data used to make Figure 6-figure supplement 1.

